# Active Motor Preparation Drives Corticospinal Facilitation during Interception Planning

**DOI:** 10.64898/2026.01.13.699289

**Authors:** Justin R. McCurdy, Brendan Jarvis, Emma Whitman, Krithi Ariga, Asher Khan, Deborah A. Barany

**Affiliations:** Department of Kinesiology, University of Georgia, Athens, GA 30602; Neuroscience Program, University of Georgia, Athens, GA 30602; Department of Interdisciplinary Biomedical Sciences, University of Georgia, Athens, GA 30602; Regenerative Bioscience Center, University of Georgia, Athens, GA 30602

**Keywords:** corticospinal excitability, motor preparation, eye movements, interception planning, visuomotor control, transcranial magnetic stimulation

## Abstract

Intercepting a moving target requires integrating visual motion information with motor preparation, yet how active motor preparation and gaze strategy jointly shape corticospinal excitability during interception planning remains unclear. We applied single-pulse transcranial magnetic stimulation over primary motor cortex to probe corticospinal excitability while participants either prepared to intercept a moving target or passively viewed its motion. Targets moved at one of two speeds, and participants judged target speed relative to a reference after each trial. Across blocks, participants were instructed to either smoothly pursue the target or fixate their eyes on the interception zone. Motor-evoked potentials were elicited at baseline, after task instruction, early during target motion, or shortly before target arrival. Task and gaze strategy interacted to influence both perceptual judgments and corticospinal excitability. Preparing to intercept produced a robust facilitation of corticospinal excitability immediately before movement onset, regardless of gaze strategy. In contrast, corticospinal excitability during passive viewing did not differ from baseline. Smooth pursuit improved perceptual accuracy and reduced interception timing error compared with fixation but did not independently influence corticospinal facilitation. These findings demonstrate that active motor preparation is the primary determinant of the transition to a state of facilitated excitability during interception planning, whereas eye movements modulate perceptual estimates and support behavioral performance.

**New & Noteworthy:** This study dissociates the contributions of visual tracking, eye movements, and active motor preparation to corticospinal excitability during interception. Here, we show that corticospinal facilitation emerges primarily when an action is planned, whereas passive viewing remains at baseline despite identical visual input. Smooth pursuit enhanced perceptual accuracy and interception timing but had a limited effect on corticospinal excitability, highlighting active motor preparation as the key driver of corticospinal output.

## Introduction

Intercepting a moving object relies on the brain’s ability to rapidly predict a target’s trajectory and time an appropriate motor response. Visual motion cues during interception shape perceptual judgments and prime the motor system for action, supporting both spatial and temporal accuracy under variable dynamics and environmental uncertainty^1–3^. Because interception requires integrating visual motion signals with motor commands, these cues may not only inform perceptual judgments but also provide predictive information about target kinematics^4,5^. In turn, engaging the motor system can enhance motion perception^6^, suggesting a bidirectional link between visual motion perception and interception planning.

Additionally, gaze behavior plays a crucial role in visually guided interception^7^. Smooth pursuit eye movements (SPEM) stabilize foveal input by matching ocular velocity to target speed and providing continuous, high-resolution feedback^8,9^. In contrast, endpoint fixation holds gaze on an anticipated interception zone, relying on peripheral cues and internal predictive models to estimate target arrival. In scenarios with predictable motion, fixating on an interception location can be advantageous, whereas when motion is less predictable, SPEM tends to enhance interception performance^7,10,11^. Taken together, these findings indicate that interception performance emerges from the coordinated interactions of motor preparation, visual motion processing, and gaze behavior.

In humans, transcranial magnetic stimulation (TMS) has been used to non-invasively probe corticospinal excitability (CSE) across movement stages. Preparatory CSE typically exhibits a biphasic pattern, with an early inhibitory phase followed by a facilitation peak immediately before movement onset^12–15^. Previous research has identified that the extent of CSE modulation scales with task demands during action preparation, such as switching between plans to initiate, withhold, or cancel a response^16–21^. CSE is also affected by the complexity of the motor plan^22^ and modulated by visual information, such as uncertainty of motion cues^23,24^. Therefore, modulation of CSE during interception planning may reflect contributions from both active motor preparation and the integration of task-relevant visual cues.

Even in the absence of overt movement preparation of the targeted effector, visual input and task demands can modulate CSE. Classic studies using action observation paradigms have found that CSE increases when watching the actions of others^25–27^. This facilitation is enhanced when the visual stimuli are behaviorally relevant^28^, paired with imagery^29,30^, or specific to motion kinematics^31^. The type of visual feedback of one’s own actions (e.g., movement of the opposite limb) can also modulate CSE in an effector-specific pattern^32,33^. These results support the hypothesis that CSE reflects the priming of the motor system for simulation of goal-directed action^34^.

Some of the visual-related modulation of CSE in the intrinsic hand muscles during action observation may be attributed to differences in eye movements across conditions. For example, enhanced CSE is associated with more fixations on goal- relevant information^28^. Smooth pursuit eye movements can influence motor readiness by modulating long-latency reflexes in the lower limb during stance, with reflex responses to perturbations significantly larger during pursuit than during fixation^35^. In addition, smooth pursuit eye movements in the absence of visual feedback globally suppresses CSE of the hand muscles^36^, indicating that oculomotor commands can directly influence excitability. These findings suggest that gaze behavior is not merely a passive consequence of visual processing, but an active contributor to motor preparatory states^37^.

Our previous work found evidence that CSE is facilitated while preparing to intercept faster moving targets, demonstrating that visual motion kinematics can influence preparatory dynamics^5^. However, the observed differences between intercepting fast and slow targets could arise from multiple sources, including differences in visual input, differences in imposed task demands, and differences in gaze strategy. In the present study, we aimed to disentangle the respective contributions of these different factors to changes in corticospinal output during timed interceptions. We measured the effects of TMS-elicited motor-evoked potentials (MEPs) during interception planning and passive observation of moving targets. We hypothesized that active movement preparation is a necessary condition for CSE modulation, then only interception planning should elicit changes in CSE while viewing moving targets. Alternatively, if changes in visual motion information or gaze are sufficient, then we would expect a similar pattern for passive observation.

To examine the role of gaze behavior and visual input on CSE modulation, we additionally instructed participants to adopt one of two gaze strategies: continuously track the moving target via smooth pursuit (“Follow”) or to maintain fixation on the interception zone without pursuing the target (“Fixate”). Based on previous work, we hypothesized that engaging the smooth pursuit system would modulate CSE in both active preparation and passive observation task contexts.

We found late facilitation exclusively during active interception planning, with limited modulation by eye movement strategy. This pattern demonstrates that active motor preparation, rather than changing sensory input alone, drives the transition from suppression to facilitation in corticospinal circuits. These results clarify how the motor system integrates predictive visual information with preparatory commands to support accurate interception.

## Methods

### Participants

Twenty healthy volunteers (10 M, 23.7 ± 4.3 years) took part in the study. All participants were right-handed, had normal or corrected-to-normal vision, reported no history of neurological disorders, and had no contraindications to TMS. The target sample size was selected to be consistent with prior single-pulse TMS studies examining corticospinal excitability during movement preparation and timed interception (e.g., Ibáñez et al., 2020; McCurdy et al., 2025). Written informed consent was obtained from all participants, and the study was approved by the Institutional Review Board at the University of Georgia, in accordance with ethical guidelines for human research.

### Experimental Setup

Participants were seated in front of the task monitor (1920 x 1080 resolution, 60 Hz refresh rate) at a desk with their right hand resting on a digitizing trackpad (WACOM Intuous Pro Large). The experiment and stimulus presentation were controlled using PsychoPy3^38^. The trackpad was positioned so that participants could rest both arms comfortably on the desk, while placing their chin in a custom stand approximately 58 cm from the center of the screen. Participants wore earplugs for the duration of the study and a sleeve over their index finger to enhance trackpad sensitivity. Surface electromyography (EMG) activity of the FDI and ADM muscles of the participant’s right hand were recorded using a Delsys Trigno wireless system (Delsys Inc., Boston, MA, USA). EMG signals were amplified 1000x, bandpass filtered (20-450 Hz), and sampled at 5000 Hz through a data acquisition board (Micro 1401-4, Cambridge Electronic Design, Cambridge, UK) and CED Signal software. Monocular eye position was recorded at 500 Hz and monitored in real-time using a high-precision, video-based eye-tracking system (Eyelink 1000 Plus, SR Research, Ottawa, ON, Canada).

### Transcranial Magnetic Stimulation

TMS was applied through a 70-mm, figure-of-eight coil driven by a DuoMAG MP magnetic stimulator (Deymed Diagnostics sro, Hronov, Czech Republic). MEPs were elicited from the right FDI by applying TMS over the hand area of the left M1. The coil was placed tangentially on the scalp with the handle oriented towards the occiput and laterally from the midline at a 45° angle. Before each of the experimental sessions, the FDI motor hotspot (i.e., the optimal location for eliciting MEPs in the target muscle) was identified on each participant by eliciting TMS every 4 s and systemically repositioning the coil. At the beginning of each experimental session, a model of the brain was scaled to the participant’s cranial dimensions using a neuronavigational tracking system (BrainSight, Rogue Research, Montreal, Canada) to optimize and replicate coil position throughout the session. Resting motor threshold (rMT) was defined as the minimal TMS intensity required to evoke peak-to-peak MEP amplitudes of ∼50 uV in the target muscle for 5 of 10 consecutive trials^39^. The average rMT for this experiment was 41.1% (± 4.7) of the maximum stimulator output. Stimulator intensity during the task blocks was set to 115% of the participant’s rMT. TMS timings were controlled by sending TTL pulses from the task computer to the stimulator via Signal software (Cambridge Electronic Design, Cambridge, UK).

### Experimental Design and Procedure

Participants performed a combined speed discrimination and timed interception task in which they judged the speed of a moving target while either executing a predictively timed finger-swipe movement to intersect the target at a predefined interception zone (IZ) or passively viewing its motion (Fig.1). On separate experimental days, participants employed different gaze strategies: they were instructed to either smoothly track the target with their gaze (“Follow“) or maintain fixation on the interception zone (“Fixate“) until the target arrived.

**Figure 1.**
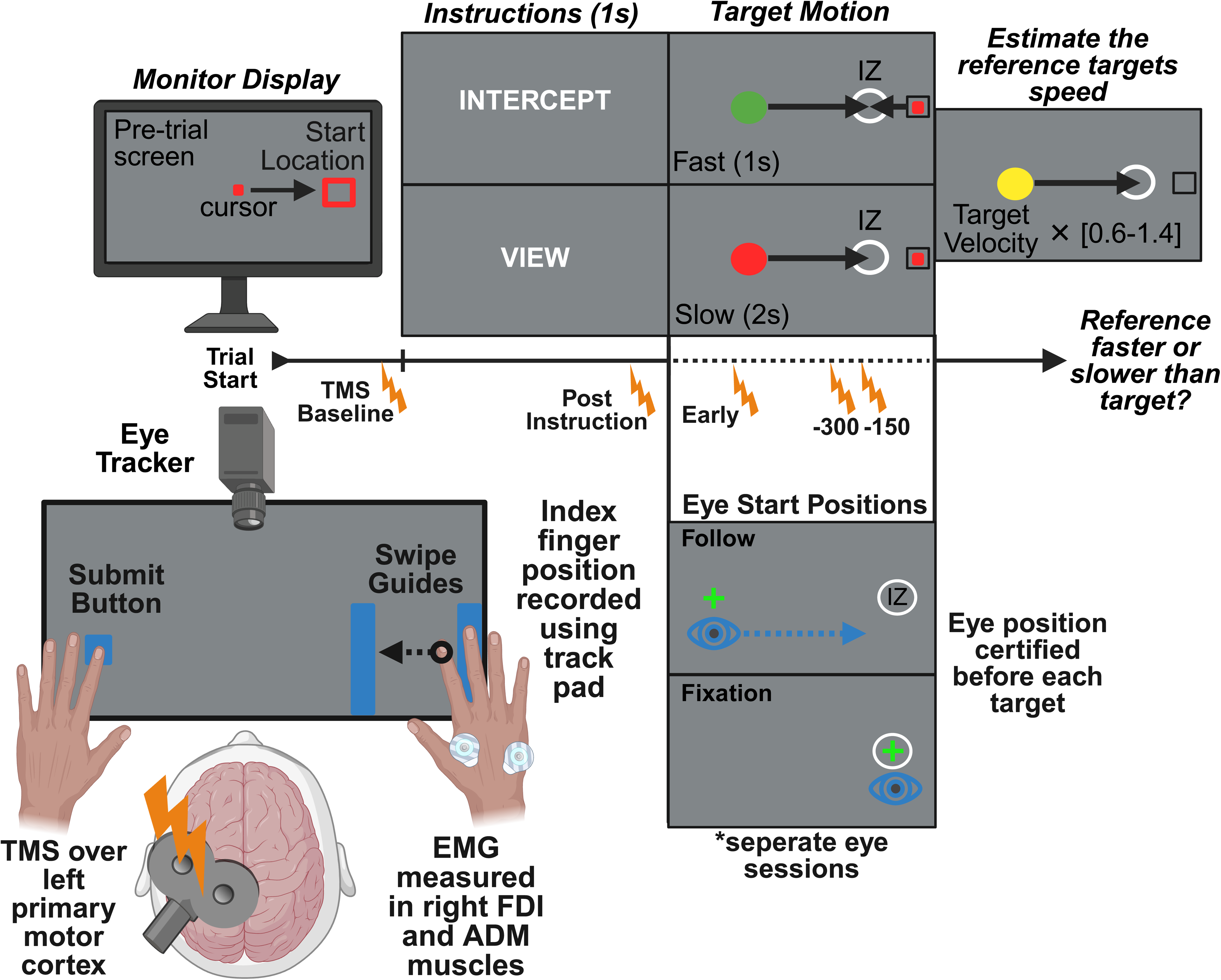
Interception task design. Participants completed an interception task requiring them to execute a timed swipe to intersect a moving target at a predefined interception zone (IZ) while following assigned eye movement strategies (Fixate or Follow). The trackpad included two tactile swipe guides (blue bars) that provided physical boundaries for the required swipe trajectory, ensuring that participants moved fully through the IZ on Intercept trials. A response button (blue circle) on the left side of the trackpad was pressed with the left hand to submit the speed discrimination judgment. During the pre trial phase, participants positioned the on screen cursor, shown as a red dot inside a black start location box, and held it there until cued to begin the trial. The target moved at one of two constant velocities (Fast or Slow), and after each trial, participants judged whether a reference target was faster or slower than the original. Eye tracking recorded gaze position, and TMS was applied over left M1 at baseline or at one of three preparation time points. EMG was recorded from the right FDI and ADM muscles.

To standardize hand movements, the trackpad included two tactile swipe guides (thin tape strips) that served as physical boundaries indicating the required start and end positions of the swipe. These guides ensured that participants consistently executed a full amplitude swipe through the IZ on every interception trial. A response button (blue circle in Fig. 1), pressed with the left hand, was used to submit the speed discrimination judgment after each trial; this functioned as a simple mouse click to confirm the participant’s selection on the continuous rating scale.

In the pre trial phase, participants moved their right index finger on the touchpad to position an on screen cursor, displayed as a red dot within a black start location box, into a fixed start position. The cursor remained inside this box until the trial began, ensuring that participants held a stable starting posture and initiated their swipe only at the appropriate time. Once the cursor remained in the start location for 100 ms, participants were cued to either prepare a timed interception movement intercept (“Intercept”) or simply view its motion (“View”). After 1 s, the instruction cue disappeared, and participants moved their eyes to the appropriate eye start location: either the target’s start position for the Follow session or the interception zone for the Fixation session, indicated by a small green fixation cross that disappeared once gaze was detected for more than 300 ms.

After the pre-trial phase, a green (Intercept trials) or red (View trials) target (circle, 0.7 cm radius) would appear 24 cm to the left of the interception zone. After a brief static delay (300 ms), the target swept across the screen at one of two constant horizontal speeds, 12 cm/s (“Slow”) or 24 cm/s (“Fast”), until it reached the interception zone. On Intercept trials, participants were instructed to abduct their right index finger to execute a timed swipe that intersected the target as it entered the interception zone.

During training blocks, the target paused briefly upon interception to provide feedback about the target’s distance from the interception zone at the moment the cursor reached it. In all subsequent blocks, the target disappeared as soon as the cursor entered the interception zone. On View trials, participants were instructed to keep their hand at rest throughout the trial.

Once the first target either stopped due to a successful interception or passed through the interception zone during View trials, participants were instructed to move their eyes back to the start position for that trial. Once gaze position was confirmed, a yellow reference target appeared and remained static for 300 ms before retracing the same horizontal motion path as the primary target, but at a different speed. The speed was drawn at random from six multipliers of the trial speed (0.6, 0.8, 0.9, 1.1, 1.2, or 1.4), making it slower or faster than the primary target (24 vs. 12 cm/s). During the training period, participants were explicitly instructed to compare the speeds of the two targets and to submit their judgment after making a decision. After watching the reference target motion, participants indicated whether it moved faster or slower than the primary target by sliding a cursor along a continuous scale and pressing a button with their left hand to submit their judgment.

Prior to the main task, participants completed repeated practice blocks consisting solely of interception trials until they reached at least 80% accuracy, defined as executing a swipe that intersected the target within 90 ms of its arrival at the interception zone. Each practice trial included immediate feedback (“Too Early,” “Too Late,” or “Perfect”). After meeting criterion, participants completed a 48 trial block with unrestricted eye movements, followed by a 48 trial eye training block in which they were instructed on the designated gaze strategy for that session. No TMS was applied during these training blocks. Both Intercept and View trials were included, and gaze behavior was continuously monitored to ensure compliance with fixation or smooth pursuit instructions. Participants then performed a no TMS behavioral baseline block to characterize performance in the absence of stimulation.

To evaluate CSE modulation during preparation, transcranial magnetic stimulation (TMS) was applied on 43 of the 49 trials in each task block under three different timing conditions: (1) a baseline stimulation delivered prior to the instruction window when no stimuli were present (3 trials per block), (2) a post-instruction stimulation administered after the response cue but before the target’s appearance (4 trials per block), and (3) trial stimulation delivered at one of three specific time points: 200 ms after target onset (“Early”), 300 ms before target arrival at the interception zone, or 150 ms before target arrival (12 trials per timepoint per block). The “Early” timing was not intended to sample the early 0–150 ms inhibitory window described in classical preparatory suppression studies^14,40–42^. Instead, the 200 ms post motion onset probe was chosen to mark the beginning of interception related processing, when participants start extracting target speed information, assessing urgency, and initiating early timing preparation. This probe allowed us to test whether suppression at motion onset varied with task goals or perceived urgency, building on prior work from our group showing target speed dependent suppression well before movement onset^5^. In the remaining 6 trials, no TMS was applied.

The order of the eye movement strategy condition for each experimental session (“Follow” or “Fixate”) was counterbalanced across participants. In each experimental session, participants performed 5 task blocks of 49 trials each, for a total of 245 trials (215 TMS trials and 30 No-TMS trials). Task instruction (Intercept vs. View), target speed (Fast vs. Slow), and TMS timing (Baseline, Post-instruction, Early, 300 ms, 150 ms) were pseudo-randomized across trials to minimize anticipation effects. Altogether, there were 15 trials performed per condition on either day (e.g., 15 Baseline-Fixate trials).

### Data Analysis

EMG recordings were analyzed offline using the MATLAB VETA toolbox ^43^. Interception performance was quantified by movement onset, movement time, and interception timing. Movement onset was identified as the first timepoint EMG activity in the FDI muscle exceeded 3 standard deviations of the mean of the rectified signal for the entire trial epoch and was >0.5 mV, relative to the target’s arrival time. Movement time was defined as the duration from EMG-derived movement onset to the time at which the interception zone was reached. Interception timing error was defined as the absolute timing error between the time the target and finger position entered the interception zone.

To quantify speed perception, participants’ judgments on the continuous scale were converted to binary responses indicating whether the reference target was judged as faster or slower than the primary (test) target. We defined RefScale as the reference speed divided by the test speed and grouped trials by RefScale. For each subject condition, the proportion of “reference faster” responses were fit with a sigmoid two-parameter logistic psychometric function using nonlinear least-squares fitting in Python (*curve_fit* function from *scipy.optimize* package), from which we extracted the point of subjective equality (PSE). PSE values reported in RefScale units and expressed as percent speed difference for plotting. Because RefScale is the reference speed divided by the test speed, a PSE greater than 1 means the reference had to be faster than the test for the two to be judged equal, which indicates the primary test target was perceived as faster than it actually was (overestimation). A PSE less than 1 means the primary test target was judged as slower than it actually was (underestimation).

For Follow eye movements, smooth pursuit eye movements were identified when gaze and object locations remained continuously within a foveal visual radius of ∼2° visual angle from the moving target, consistent with established criteria for pursuit detection ^8,9^. Saccades were detected when instantaneous eye velocity exceeded 30°/s or acceleration exceeded 8,000 °/s^2^, indicating rapid shifts in eye position ^44^. SPEM gain, a measure of SPEM quality, was calculated by dividing gaze angular velocity by target angular velocity over the middle 40% of each target motion period. A SPEM gain of 1 indicates ideal target tracking; less than 1 indicates the eye lagged the target; greater than 1 indicates the eye led the target.

CSE was indexed by the TMS-elicited MEP peak-to-peak amplitude. The root mean square (RMS) value of the background EMG activity was calculated for the 300 ms before the TMS artifact. Trials were rejected from analysis when the background RMS was ±2 standard deviations from the participant’s average RMS or when EMG activity preceded the MEP to account for changes in MEP amplitude due to fluctuations in background EMG activity^45^. The peak-to-peak amplitude of the EMG signal was calculated during the 20-50 ms window after TMS (i.e., the normal range at which an MEP is expected to occur). MEP amplitudes were normalized to the participants average session-specific within-task baseline.

### Statistical Analysis

All primary analyses were specified as *a priori* based on the experimental design. Statistical analyses were performed in R (v4.2.1+) using the *lme4*, *lmerTest*, and *emmeans* packages, with a two-tailed α level of 0.05. Prior to modeling, each outcome measure was screened for normality (Kolmogorov–Smirnov) and variance homogeneity (Levene’s test). Given our within-subject design, we fit linear mixed-effects models (LMMs) using restricted maximum likelihood (REML). Fixed effects were assessed via Type III ANOVA using Satterthwaite’s approximation for denominator degrees of freedom, and random effects structures were selected based on likelihood ratio tests; for all behavioral outcomes, models with participant specific intercepts provided the maximal non singular structure and were therefore retained. Post hoc pairwise comparisons were conducted on estimated marginal means using Tukey adjustment, which is appropriate for all pairwise contrasts among factor levels in mixed effects models and preserves the model based covariance structure. To evaluate evidence for or against null effects, we conducted Bayesian paired samples tests using default JZS Bayes factors (r = 0.707) implemented in the BayesFactor package^46^. Bayes factors were interpreted using conventional thresholds, with BF_10_ < 1/3 (i.e., BF_01_ > 3) indicating evidence for the null model, BF_10_ > 3 indicating evidence for the alternative, and values between 1/3 and 3 considered inconclusive.

Smooth pursuit gain was modeled with Target Speed (Fast vs. Slow) and Task Goal (Intercept vs. View), and their interaction. The model included participant-level random intercepts and random slopes for Target Speed. Perceptual thresholds (PSEs) were analyzed using a linear mixed effects model with fixed effects of Target Speed, Eye Strategy, Task, and their interactions. The random effects structure included participant specific intercepts. More complex random effects structures (e.g., by strategy slopes) resulted in singular fits and were therefore not retained.

Interception kinematics (i.e., movement onset latency, movement time, and normalized interception timing), were each analyzed with LMMs including fixed effects of Eye Strategy and TMS Timing (NoTMS, Early, −300 ms, −150 ms) and participant specific random intercepts.

Corticospinal excitability was examined in two stages: (1) pre-trial instruction effects were assessed with an LMM predicting in-session normalized MEP amplitude at 200 ms post-cue from Eye Strategy and Instruction, covarying baseline EMG (RMS), and followed by Holm-corrected one-sample t-tests against baseline for each Eye × Instruction cell; (2) in-trial stimulation effects were tested with an LMM predicting in-session normalized MEP amplitude including Task, Eye Strategy, and TMS Timing, covarying pre-stimulation EMG (RMS) and modeling by-factor random slopes, with subsequent one-sample t-tests comparing each Task × Eye Strategy × TMS condition to baseline.

## Results

### Task and gaze strategy influenced judgments of target speed

We first assessed how movement preparation and eye movement strategy influenced perceived target speed. Figure 2A,B shows the group averaged psychometric functions for Fast and Slow targets. As expected, the functions differ in slope, though this likely reflects the scaling of the reference stimulus rather than true differences in sensitivity. Figure 2C,D plots the resulting points of subjective equality (PSEs). We define RefScale as the reference speed divided by the test speed, and we report PSE − 1 so that zero corresponds to perceptual equality. Negative PSE − 1 values indicate that the test was perceived as slower than it actually was, and positive PSE − 1 values indicate that the test was perceived as faster than it actually was.

**Figure 2.**
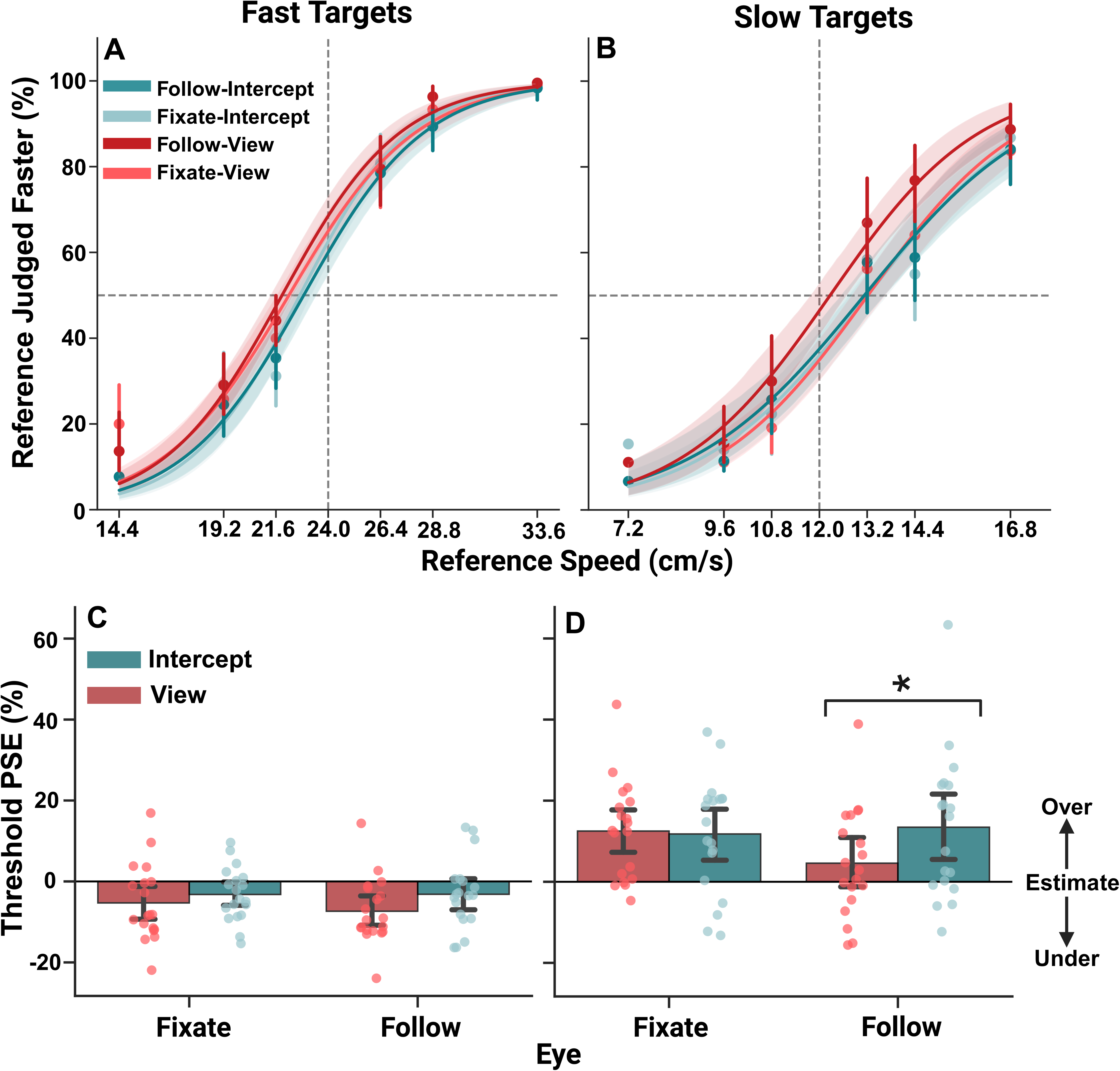
Overestimation of slow targets and underestimation of fast targets. (A– B) Group average psychometric functions for Fast (A) and Slow (B) targets under fixation (dashed lines) and smooth pursuit (solid lines) in both View and Intercept conditions. The reference speeds ranged from 0.6 to 1.4 times the primary target speed. (C–D) Group averaged points of subjective equality (PSE) for Fast (C) and Slow (D) targets, with 95% confidence intervals. Panels C–D plot PSE − 1 so that zero corresponds to perceptual equality. Negative values indicate the test was perceived as slower than the reference (underestimation); positive values indicate the test was perceived as faster than the reference (overestimation). *p < .05 between conditions (Holm-corrected).

Group means of PSE − 1 show a clear velocity effect. For Fast targets the mean PSE − 1 was −0.0475 (SEM = 0.0095, n = 76), corresponding to a −4.75 percent shift. For Slow targets the mean PSE − 1 was 0.1058 (SEM = 0.0170, n = 76), corresponding to a +10.58 percent shift. These descriptive statistics indicate that perceived speed differed systematically with target velocity, with Fast targets shifted toward negative values and Slow targets shifted toward positive values, a pattern consistent with regression toward the mean in magnitude estimates.

A repeated measures ANOVA confirmed a strong main effect of target speed, *F_(1, 32.3)_* = 24.38, *p* < .001, partial η^2^ = .43. Bayesian tests supported this pattern, showing strong evidence for a velocity effect across all Eye × Task combinations (BF_10_ range = 16.7 - 1315.5), and follow-up t-tests confirmed Slow–Fast differences in every condition (all *p*’s < .003).

The main effect of Task was also significant, *F_(1, 18.72)_* = 7.29, *p* = .014, partial η^2^ = .28, indicating that participants judged the primary target as moving faster when preparing an interception. However, Bayes factors provided at most weak evidence for the null, with most values falling in the inconclusive range (BF_10_ range = 0.32 - 0.99), and post hoc comparisons showed no reliable Intercept–View differences within any Eye × Velocity combination (all *p*’s > .13). There was likewise no main effect of gaze strategy, *F_(1, 18.88)_* = 2.06, *p* = .167, partial η^2^ = .10, with Bayesian tests again showing at most borderline support for the null, with evidence otherwise inconclusive (BF_10_ range = 0.32 - 1.27).

Importantly, the Task × Eye interaction was significant, *F_(1, 71.94)_* = 5.91, *p* = 0.018, partial η^2^ = .08. Consistent with this interaction, PSEs were more positive in Intercept than View trials when participants followed the target (Holm-corrected *p* < .007), whereas no difference emerged during fixation (*p* = 1.00). Thus, the largest PSE shifts occurred when interception was paired with smooth pursuit. Because PSE reflects both perceptual and decisional components^47,48^, these shifts cannot be attributed solely to changes in sensory encoding. Instead, the interaction indicates that the combined influence of gaze strategy and task goal systematically altered the judgment process underlying speed reports^49^. Overall, speed judgments were shaped primarily by target velocity, with additional modulation by task goal during smooth pursuit.

### Smooth pursuit eye movements improved interceptive timing

TMS timing significantly affected movement onset, *F_(3, 2990.2_)* = 93.14, *p* < .001, partial η^2^ = .085 (Fig. 3A). Consistent with this main effect, stimulation delivered 150 ms before target arrival produced markedly delayed movement initiation relative to all other TMS timings (β range = 30.7 - 38.1 ms; all *p* < 0.001), indicating disruption of anticipatory preparation. when stimulation occurred closest to movement onset. Bayesian comparisons strongly supported this pattern, providing evidence for delayed initiation at −150 ms relative to −300 ms, Early, and No TMS stimulation (BF_10_ = 7.98 × 10 - 5.06 × 10). In contrast, neither Early or −300 ms stimulation differed from the No TMS condition (all *p* > .05), and Bayesian tests indicating evidence in favor of the null (BF_10_ = 0.18 - 0.30). Eye movement strategy did not independently influence movement onset, *F_(1, 2992.7)_* = 0.01, *p* = .922, partial η^2^ < .001, and the Eye × TMS interaction was similarly nonsignificant, *F_(3, 2989.7)_* = 0.25, *p* = .862, partial η^2^ < .001.

**Figure 3.**
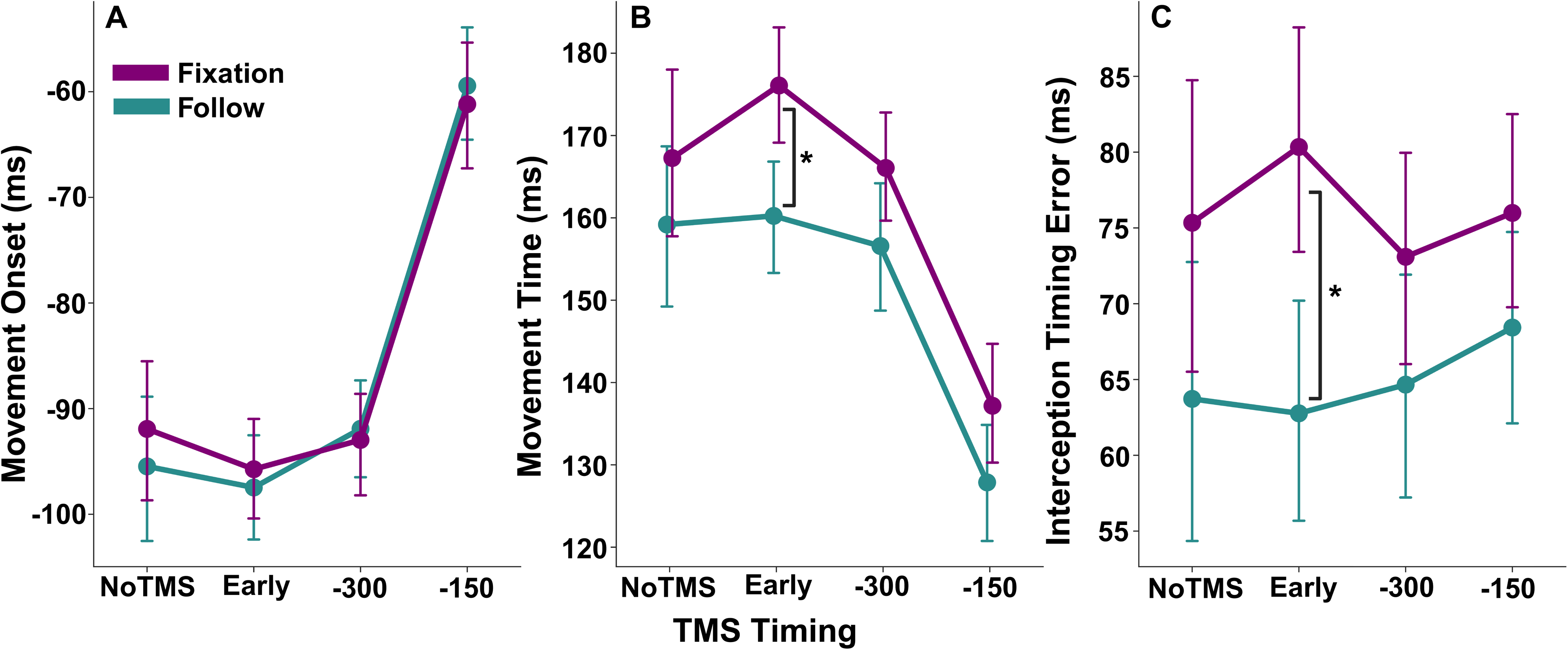
Smooth pursuit eye movements improved interception timing. (A) Movement onset relative to target arrival, showing later initiation for -150ms compared to NoTMS. (B) Movement time, indicating shortened duration for late simulation (-150). (C) Interception timing error, demonstrating improved accuracy when following the target with gaze. Error bars represent 95% confidence intervals. *p < 0.05 between conditions (Tukey-adjusted).

Bayesian simple effects tests ranged from weak evidence for the null to inconclusive evidence (BF_10_ = 0.24 - 0.45). Fixed effect estimates for Eye and all interaction terms were small (β range = −3.88 to 1.81 ms) with 95% CIs spanning zero, indicating that movement initiation was modulated by the timing of TMS, independent of gaze strategy.

Movement time was strongly modulated by TMS timing, *F_(3, 2989.2)_* = 68.46, *p* < .001, partial η^2^ = .064, with later stimulation producing progressively faster movements (Fig. 3B). Pairwise contrasts showed that −150 ms stimulation yielded substantially shorter movement times than all other timings (all p < .001), a pattern supported by Bayesian evidence for differences between −150 ms and both −300 ms and Early stimulation (BF_10_ = 3.9 × 10 - 1.7 × 10) and strong evidence relative to No TMS (BF_10_ = 2.65 × 10). Earlier stimulation times did not differ from one another (BF_10_ = 0.20 - 0.72). Eye strategy also influenced movement time, with Follow trials producing faster movements overall, *F_(1, 2989.6)_*= 29.16, *p* < .001, partial η^2^ = .010, and modest Bayesian support (BF_10_ = 1.63). A Eye × TMS interaction was not significant, F(3, 2989.1) = 0.63, *p* = .599, partial η^2^ < .001, and Bayesian simple effects tests favored the null or were inconclusiveacross TMS levels (BF_10_ = 0.27–1.50). Fixed effect estimates for interaction terms were small (β = −5.85 to 0.83 ms), indicating that although smooth pursuit generally sped up movement execution, this benefit did not depend on when TMS was delivered.

Eye movement strategy significantly influenced interception timing, *F_(1, 2990.1)_*= 24.70, *p* < .001, partial η^2^ = .008 (Fig. 3C). Consistent with this main effect, the Follow strategy was associated with reduced timing error relative to Fixate (β = −6.40 ms, 95% CI [−13.8, 1.0]), indicating more precise interception when participants tracked the moving target. Bayesian analysis provided strong support for this Eye effect (BF_10_ = 108.7). In contrast, the effect of TMS timing was small, *F_(3, 2989.4)_* = 2.82, *p* = .038, partial η^2^ = .003, and Bayesian pairwise tests favored the null for all TMS comparisons (BF_10_ = 0.18 - 0.28), indicating no reliable influence of stimulation timing on interception accuracy. The Eye × TMS interaction was also not significant, *F_(3, 2989.3)_* = 1.31, *p* = .270, partial η^2^ < .001, with Bayesian simple effects tests ranging from evidence favoring the null to inconclusive evidence across most TMS levels (BF_10_ = 0.27–0.56), except for modest evidence of an Eye effect during Early stimulation (BF_10_ = 14.26). Fixed effect estimates for TMS and interaction terms were small (β ranges: TMS = −7.18 to 0.51 ms; interaction = −8.58 to −0.27 ms) with confidence intervals spanning zero, indicating that interception accuracy was primarily shaped by gaze strategy rather than stimulation timing.

To test whether perceptual biases predicted interception timing, we conducted an exploratory trial-level correlation between perceptual bias and signed timing error on interception trials, excluding trials with signed timing errors greater than ±300 ms. Perceptual bias was not reliably related to signed timing error (*r* = −0.022, *p* = 0.069; BF_10_ = 0.078).

### Speed-dependent smooth pursuit gain during motion perception and interception

To assess whether movement preparation influenced smooth pursuit quality, we quantified SPEM gain during Follow blocks (Fig. 4). Target speed strongly affected pursuit gain, *F_(1, 17.8)_* = 68.94, *p* < .001, partial η^2^ = .79, with lower gain for fast targets (β = −0.0085, 95% CI [−0.0105, −0.0065]). Bayesian tests showed the same pattern, providing evidence for a speed effect in both Intercept (BF_10_ = 4.7 × 10^2^) and View trials (BF_10_ = 1.8 × 10¹^2^). In contrast, Task did not meaningfully change pursuit gain, *F_(1, 17.7)_* = 2.28, *p* = .149, partial η^2^ = .11, with Bayes factors showing inconclusive evidence for slow targets (BF_10_ = 0.88) and evidence for the null for fast targets (BF_10_ = 0.094).

**Figure 4.**
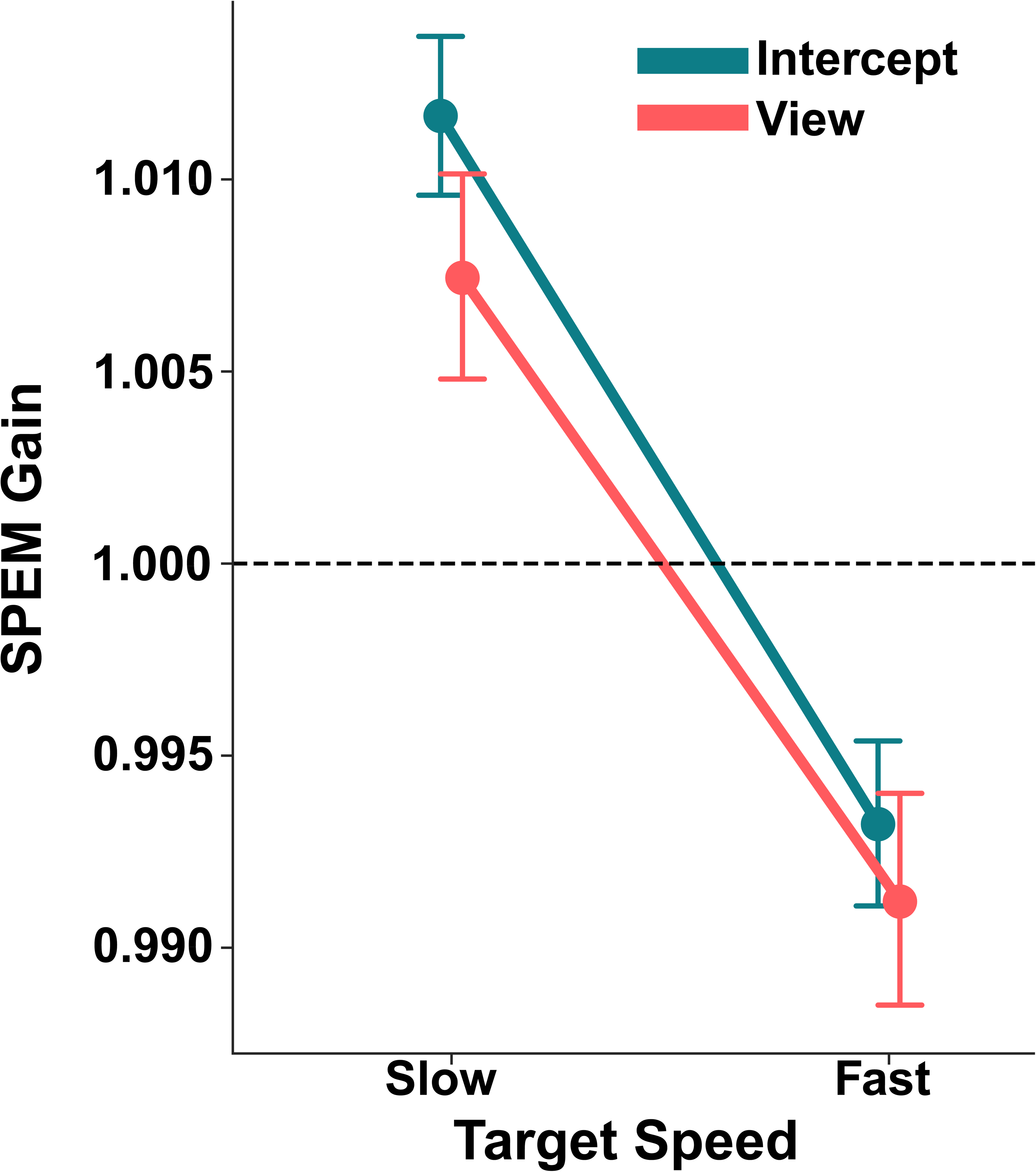
Smooth pursuit eye movement gain similar during interception and passive viewing. Eye movement velocity exceeded target velocity for slow targets and was below target velocity for fast targets for both task goals (Intercept and View). Error bars represent 95% confidence intervals.

The Speed × Task interaction was also nonsignificant, *F_(1, 3763.1)_* = 1.49, *p* = .222, partial η^2^ < .001. Random effects estimates indicated modest participant variability (SD = 0.020) relative to the residual (SD = 0.0335). Together, these results demonstrate that smooth pursuit gain was driven primarily by target speed, with highly similar gain profiles across interception and passive viewing contexts.

### Late corticospinal facilitation during interception but not passive viewing of moving targets

To evaluate how movement preparation and gaze behavior jointly shape corticospinal excitability, we analyzed FDI MEP amplitudes at three TMS latencies (Early, −300 ms, −150 ms) across gaze strategies (Fixate vs. Follow) and task goals (Intercept vs. View). MEP amplitudes were normalized to each participant’s session specific baseline, ensuring that CSE changes were evaluated relative to that session’s resting excitability.

A linear mixed effects model revealed significant main effects of Task (Intercept > View; *F_(1, 22.4)_*= 6.66, *p* = 0.017) and TMS timing (*F_(2, 40.2)_* = 5.73, *p* = 0.006), whereas gaze strategy showed no main effect (*F_(1, 24.0)_*= 0.89, *p* = 0.354). Pre TMS EMG (RMS) remained a strong positive predictor of MEP amplitude (*F_(1, 5045)_* = 39.26, *p* < 0.0001).

As shown in Figure 5, there was a significant Task × TMS timing interaction (*F_(2, 6658.0_*) = 95.30, *p* < 0.001), indicating that CSE modulation was tightly linked to movement preparation. Follow up tests using estimated marginal means showed that at −150 ms, CSE was significantly elevated during interception relative to baseline (Fixate: *t_(95.6)_*= 6.05, *p* < 0.0001; Follow: *t_(95.1)_* = 3.59, *p* = 0.0005). Bayes factor analyses supported these results, with both Fixate and Follow interception trials at −150 ms showing very strong evidence for facilitation (BF_10_ = 2.8 × 10 and 5.7 × 10). In contrast, passive viewing showed no deviation from baseline at the same latency (Fixate: *p* = 0.873; Follow: *p* = 0.559), supported by Bayes factors indicating strong evidence for the null (BF_10_ = 0.05 - 0.07). Participant-level means showed that late CSE facilitation during interception was broadly present across participants, with normalized MEPs at -150 ms exceeding baseline in 15/20 Intercept-Fixate and 15/20 Intercept- Follow participants (Supplementary Fig. 1, 10.6084/m9.figshare.32961248).

**Figure 5.**
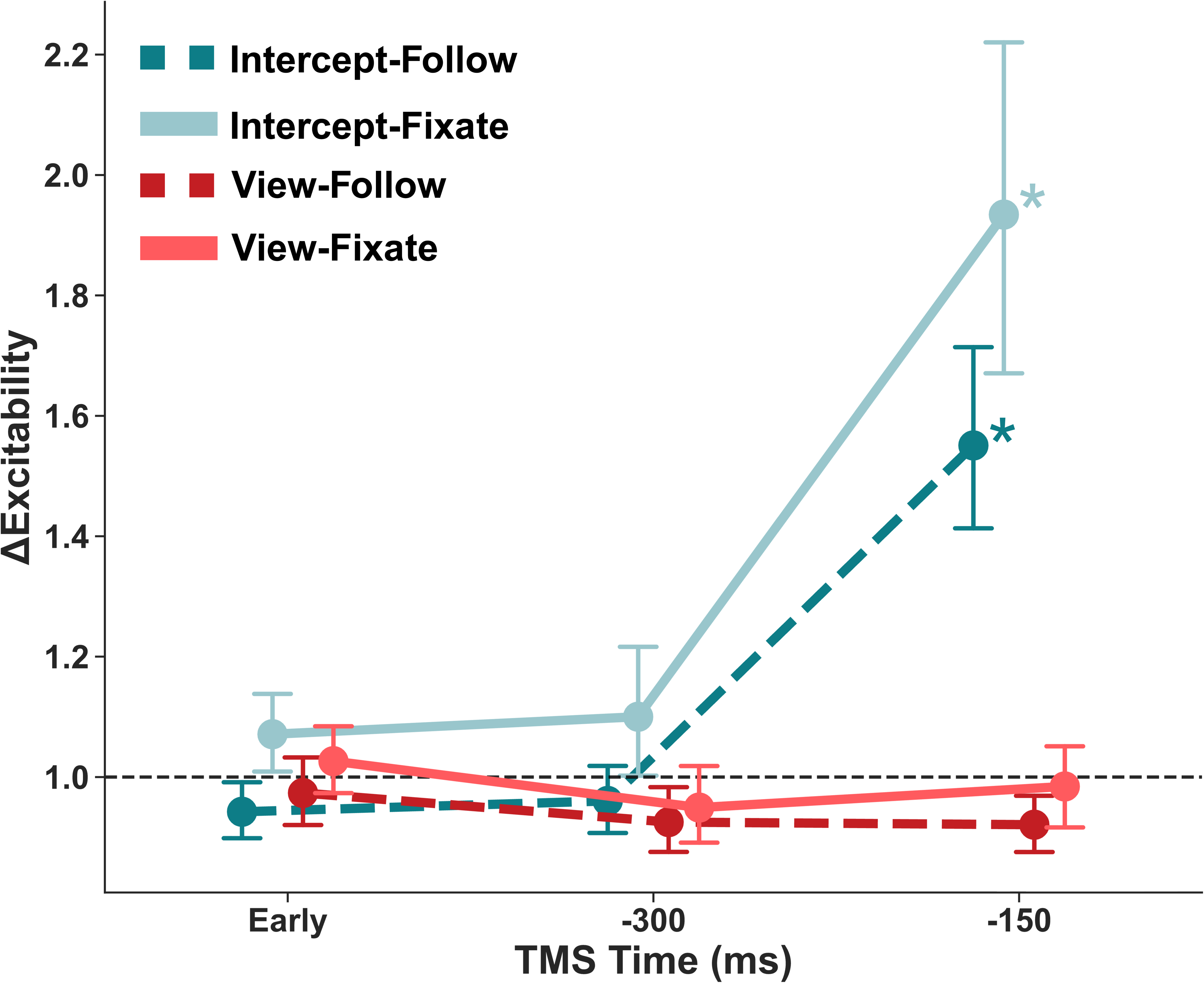
Preparation-driven facilitation of corticospinal excitability. Average normalized motor-evoked potentials for the FDI muscle are shown across three TMS time points (Early, -300 ms, and -150 ms) for each combination of eye movement strategy (Follow vs. Fixate) and task goal (Intercept vs. View). CSE increased significantly as target arrival approached the interception zone, with the strongest facilitation at -150 ms during interception trials (Follow-Intercept and Fixate-Intercept). In contrast, CSE remained near baseline during passive viewing trials (Follow-View and Fixate-View). Error bars represent 95% confidence intervals. *p < 0.05 relative to zero (Tukey-adjusted).

Across all Early and −300 ms conditions, neither Intercept nor View trials differed from baseline (all *p* > 0.05). Bayesian tests reinforced this absence of early modulation: Early and −300 ms timepoints consistently favored the null hypothesis in both tasks (BF_10_ = 0.07 - 0.38 for Fixate; 0.12 - 0.97 for Follow). The only exception was a modest deviation in View-Follow at −150 ms (BF_10_ = 5.8), but this effect was small and reflected a slight shift toward suppression.

A paired t-test showed a trend toward a baseline MEP difference between sessions (*p* = .063, Cohen’s d = 0.44), though the Bayesian paired test showed inconclusive evidence (BF_10_ = 1.156). Raw baseline MEPs were slightly higher on Follow (2.94 mV) than Fixate (2.42 mV) days, suggesting that smooth pursuit engagement may elevate baseline corticospinal excitability. However, session specific normalization removed this difference. After normalization, the magnitude of late facilitation at −150 ms did not differ between Fixate and Follow conditions (all p > 0.05; BF_10_ < 1), and the three way interaction among task goal, gaze strategy, and TMS timing was not significant (*F_(2, 6649.9)_* = 2.16, *p* = 0.116). Bayes factors likewise showed comparable evidence patterns across gaze conditions, indicating that gaze related baseline differences do not account for the interception related increase in CSE.

### Task response instruction does not significantly alter corticospinal excitability

To determine whether early task instruction modulates corticospinal excitability (CSE), we analyzed baseline normalized MEP amplitudes from TMS pulses delivered 200 ms after the instruction cue but before target motion onset. The linear mixed model revealed no main effect of eye movement strategy, *F_(1, 18.87)_* = 1.33, *p* = .263, partial η^2^ = .002, and no interaction between eye movement and task instruction, *F_(1, 749.08)_* = 0.002, *p* = .961, partial η^2^ < .001. The main effect of task instruction showed only a weak trend toward lower corticospinal excitability after the instruction to view, *F_(1, 749.05)_* = 3.20, *p* = .074, partial η^2^ = .006 (Fig. 6). As expected, pre MEP RMS strongly predicted MEP amplitude, *F_(1, 579.01)_* = 30.58, *p* < .001, partial η^2^ = .050.

**Figure 6.**
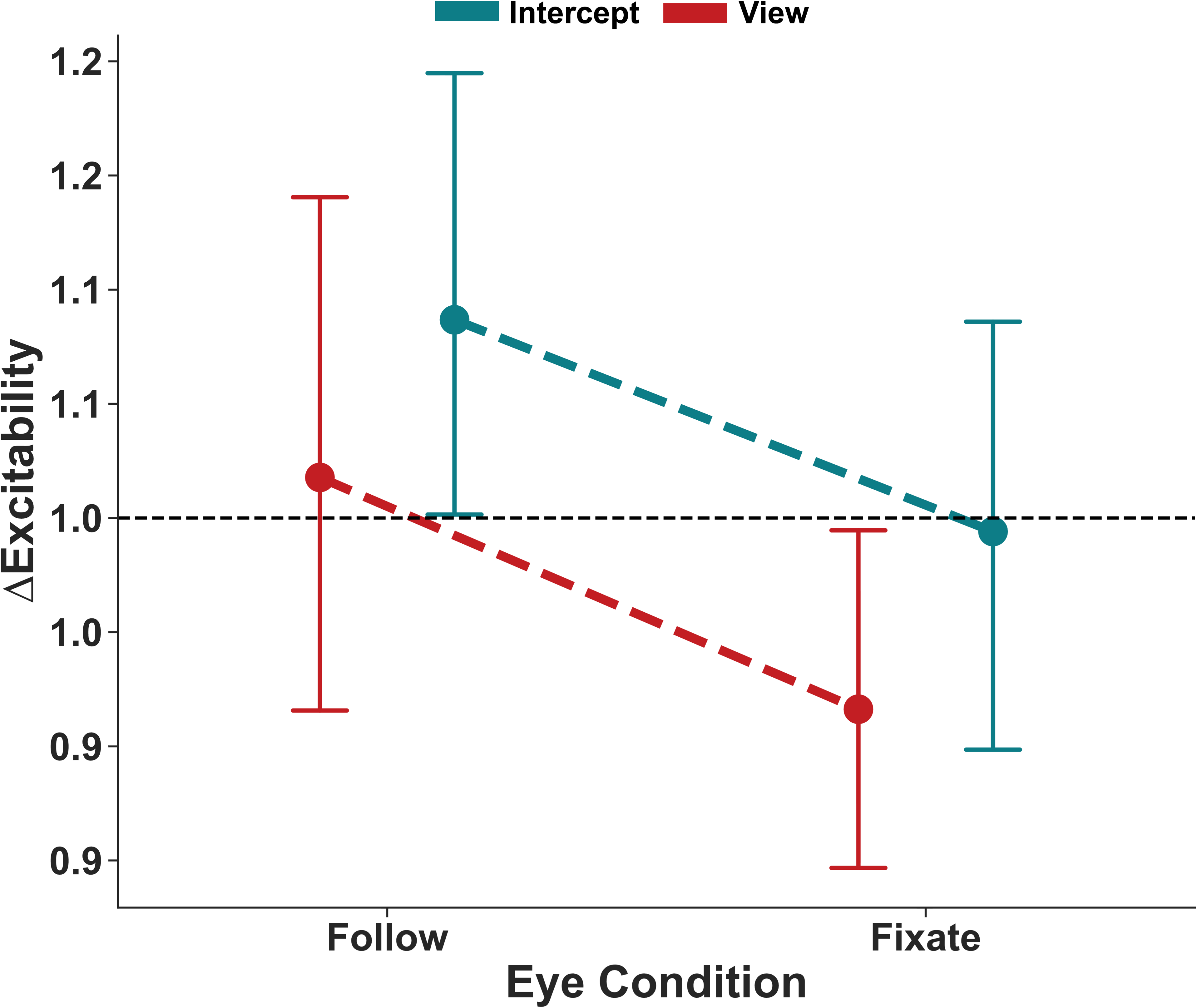
Task instruction did not significantly affect corticospinal excitability. CSE normalized to baseline for the FDI muscle are shown for each eye movement condition (Follow vs. Fixate) across task instruction conditions (View vs. Intercept). There were no significant differences in CSE between eye movement and task instruction conditions. Error bars represent 95% confidence intervals.

Estimated marginal means were close to baseline across all conditions (Intercept-Fixate: 0.998; View-Fixate: 0.919; Intercept-Follow: 1.110; View-Follow: 1.027), and none differed significantly from baseline after Holm correction (all p > .16). Bayesian one sample tests provided convergent evidence favoring the null, with Bayes factors indicating support for no deviation from baseline in the Intercept Fixate and View Follow conditions (BF_10_ = 0.08 for both), and inconclusive evidence for Intercept Follow (BF_10_ = 0.45) and View Fixate (BF_10_ = 0.72).These results indicate that task instruction did not reliably modulate CSE, and any differences between instruction or eye movement conditions were small (partial η^2^ < .01). This suggests that the initial cue to intercept or view the upcoming target does not produce robust preparatory changes in corticospinal excitability.

## Discussion

In this work, we measured TMS-elicited corticospinal excitability, eye-tracking, perceptual judgments, and movement kinematics to probe how gaze behavior and movement preparation shape motion perception, interception accuracy, and the state of the motor system. We demonstrate that movement preparation, rather than differences in visual motion processing or the instructed eye movement strategy (smoothly tracking the target vs. maintaining fixation on the interception zone), is the primary driver of late-stage facilitation in corticospinal circuits. In contrast, passive observation of a moving target resulted in suppression or no change in corticospinal excitability.

Additionally, smooth pursuit eye movements enhanced temporal accuracy and altered perception of target speed during interception. Together, these results suggest that CSE modulation reflects coordination of perceptual, oculomotor, and motor preparatory signals for precise interceptive timing.

Performance on the speed discrimination task was generally high across response contexts, consistent with classic findings that motion-sensitive cortical areas accurately encode stimulus velocity^50,51^ Across conditions, participants tended to overestimate slower targets and underestimate faster targets, consistent with regression-to-the-mean biases in speed estimation across perceptual judgment and timed response tasks^52–55^. Introducing an interceptive goal shifted judgments in the direction of overestimating the primary target’s speed, particularly when participants smoothly pursued the target. This pattern aligns with work showing that velocity judgments and interceptive actions rely on overlapping visual and oculomotor information, even if task demands may alter how that information is weighted^56,57^.

However, because the psychophysical task was not designed to dissociate perceptual from decisional biases, these shifts may reflect changes in perceptual encoding, decision criteria, memory comparison processes, dual-task demands, or other post-perceptual influences^58–60^. Thus, the observed bias should be interpreted as a change in the overall judgment process rather than definitive evidence for altered sensory representations. Consistent with this cautious interpretation, an exploratory trial-level analysis showed that the speed judgment bias did not reliably predict signed interception timing error, suggesting that trial-by-trial variation in speed judgments do not directly explain variation in interception timing.

The interaction with gaze strategy, specifically whether participants tracked the target or maintained fixation on the interception zone, further suggests that smooth pursuit modulates the speed judgment process during interception. Although the task-by-gaze interaction was most evident for slow targets, fast targets showed a qualitatively similar shift relative to their overall tendency toward underestimation. The stronger effect for slow targets may reflect their longer viewing and preparation interval, allowing gaze strategy and action preparation to exert greater influence on speed judgments, though the design was not optimized to isolate the source of this speed-specific pattern. Pursuit introduces catch up saccades and efference copy signals that may influence comparisons between the primary and reference stimuli^7,61^. Differences in the timing or frequency of catch up saccades across tasks may therefore have contributed to the larger PSE shifts observed during pursuit-based interception^57^. This pattern aligns with work showing that motion judgments and interception can rely on overlapping visual and oculomotor sources ^56,57^. In sum, our results suggest that speed judgments during interception are not a passive readout of retinal input but is dynamically shaped by gaze behavior and the brain’s preparatory state. These findings highlight the tight integration of perceptual, oculomotor, and motor planning processes that support accurate movement.

When participants planned to intercept a moving target, CSE exhibited the expected late-stage facilitation that peaked immediately before movement onset. In contrast, there was no CSE facilitation when passively viewing – instead, CSE remained at baseline (“Follow” blocks) or was suppressed relative to baseline (“Fixation”) blocks. This finding suggests that sensory observation of object motion alone is insufficient to drive corticospinal up-regulation^62^, consistent with predictive control models of anticipatory visuomotor commands^63,64^.

The delayed movement onset following TMS at −150 ms is consistent with stimulation occurring late in motor preparation. This pulse was timed relative to the target’s expected arrival at the interception zone rather than to each participant’s movement onset. Because the swipe movement required additional time to execute, stimulation at −150 ms likely occurred near the period when the movement command was being released. Elevated MEP amplitudes at this time point therefore reflect late-stage corticospinal readiness, whereas the delay in movement onset is consistent with prior work showing that TMS delivered close to movement initiation can interrupt the developing motor command and produce a silent-period-like delay in action initiation ^5,14,15,41^. In addition, nonspecific auditory or somatosensory consequences of TMS may have contributed to transient response inhibition, similar to sensory-triggered inhibition reported in the oculomotor system^65,66^.

Contrary to classic reports of an initial inhibitory phase^42,67^, and our previous study ^5^, we did not observe a reliable early suppression during interception planning. This lack of an early trough may reflect our dual-task design: while preparing the motor response, participants simultaneously encoded target speed for a subsequent discrimination test. Maintaining readiness for both perceptual and motor demands could elevate baseline excitability or attenuate transient inhibition, effectively masking a brief early suppression. This interpretation aligns with dual task and attentional load findings showing that concurrent cognitive-motor demands can reduce intracortical inhibition or increase corticospinal excitability relative to single task conditions^68–70^.

Additionally, our earliest TMS probe at 200 ms post-cue may have missed an earlier inhibitory dip occurring immediately after the instruction cue^71,72^. To fully characteriz these early dynamics, future work should incorporate denser temporal sampling around motion onset to determine whether a short lived inhibitory phase reliably precedes the later facilitation observed here.

We found a limited effect of gaze strategy on corticospinal modulation. Previous work has shown that eye movements can suppress or facilitate corticospinal excitability of the hand muscle, even when the hand is at rest^36,73,74^. In our previous study, eye movements were unconstrained, and participants likely used a combination of smooth pursuits and saccades to track the target^75^. In the present study, we explicitly instructed participants to either fixate on the interception zone or smoothly track the target and used an eye-tracker to monitor eye movements in real-time. Despite differences in interceptive performance, we observed a similar pattern of suppression and late facilitation for both fixation and smooth pursuit conditions during interception. During passive viewing, we found evidence for suppression relative to baseline for the fixation condition only – this may be related to inhibiting eye movements during target motion^76–78^. In contrast, when the oculomotor system is actively engaged, particularly during smooth pursuit, it recruits cortical and subcortical motor networks that overlap with those involved in limb movement planning^79,80^. Functional imaging and electrophysiological studies suggest that pursuit-related activity in frontal and supplementary eye fields can modulate motor cortical output, potentially priming the motor system for coordinated action^8,81^.

In our previous study, we observed evidence of initial suppression as early as 500 ms before movement onset^5^. This led us to wonder whether suppression may be related to processing the task goal itself, rather than only action-specific modulation.

Indeed, neuroimaging and electrophysiological studies have shown that top-down task goals can modulate neural activity in sensory and motor regions well in advance of movement, consistent with proactive control mechanisms^82,83^. To address this question, we measured CSE 200 ms after the instruction cue to intercept or view. We found a modest, non-significant trend toward larger MEPs when participants were instructed to intercept versus when instructed to view. Only the Fixate-View condition showed significantly more suppression relative to baseline. This suggests that merely knowing that no eye or hand movement would follow may elicit a tonic down-regulation of corticospinal output^84,85^.

## Conclusion

The present study demonstrates that timed interception of moving targets relies on the coordinated interactions among perceptual judgments, gaze behavior, and preparatory motor signals. Nevertheless, corticospinal facilitation emerges only when an action plan is engaged, indicating that active motor preparation, rather than continuous visual tracking, ultimately drives corticospinal facilitation. Smooth pursuit sharpened interception timing but did not shift CSE from suppression to facilitation, whereas passive viewing while fixating maintained baseline CSE throughout target motion. In addition, a pre trial instruction about the task goal alone was insufficient to evoke early facilitation. Complementary speed discrimination results showed robust encoding of target speed ^6^ but underestimation of target speed when participants prepared to intercept – particularly when engaging in smooth pursuit. Together, these findings indicate that gaze aids prediction, but the transition from corticospinal suppression to facilitation is primarily governed by active motor preparation.

## Supplemental Material

Supplemental Fig. S1: 10.6084/m9.figshare.32961248

## Data Availability

Source data and code for this study are openly available at 10.6084/m9.figshare.32960918

## Acknowledgments

We thank Megha Sequeira and Ria Doshi for assistance with data collection. This work was supported by the NICHD National Center of Neuromodulation for Rehabilitation (pilot grant, P2C HD086844), the University of Georgia Mary Frances Early College of Education, and University of Georgia Office of Research. All figures were created with BioRender (https://BioRender.com/1zeqbrk).

