## Supplemental Figure 1 for "Active Motor Preparation Drives Corticospinal Facilitation during Interception Planning"

**Affiliations:**

**Manuscript Number:** JN-00017-2026

**Document Contents:**

- Supplementary Figure 1. Subject-level normalized MEP amplitudes across TMS time points for each Task × Gaze condition

**Supplementary Figure 1.** Subject-level mean normalized MEP amplitudes are shown

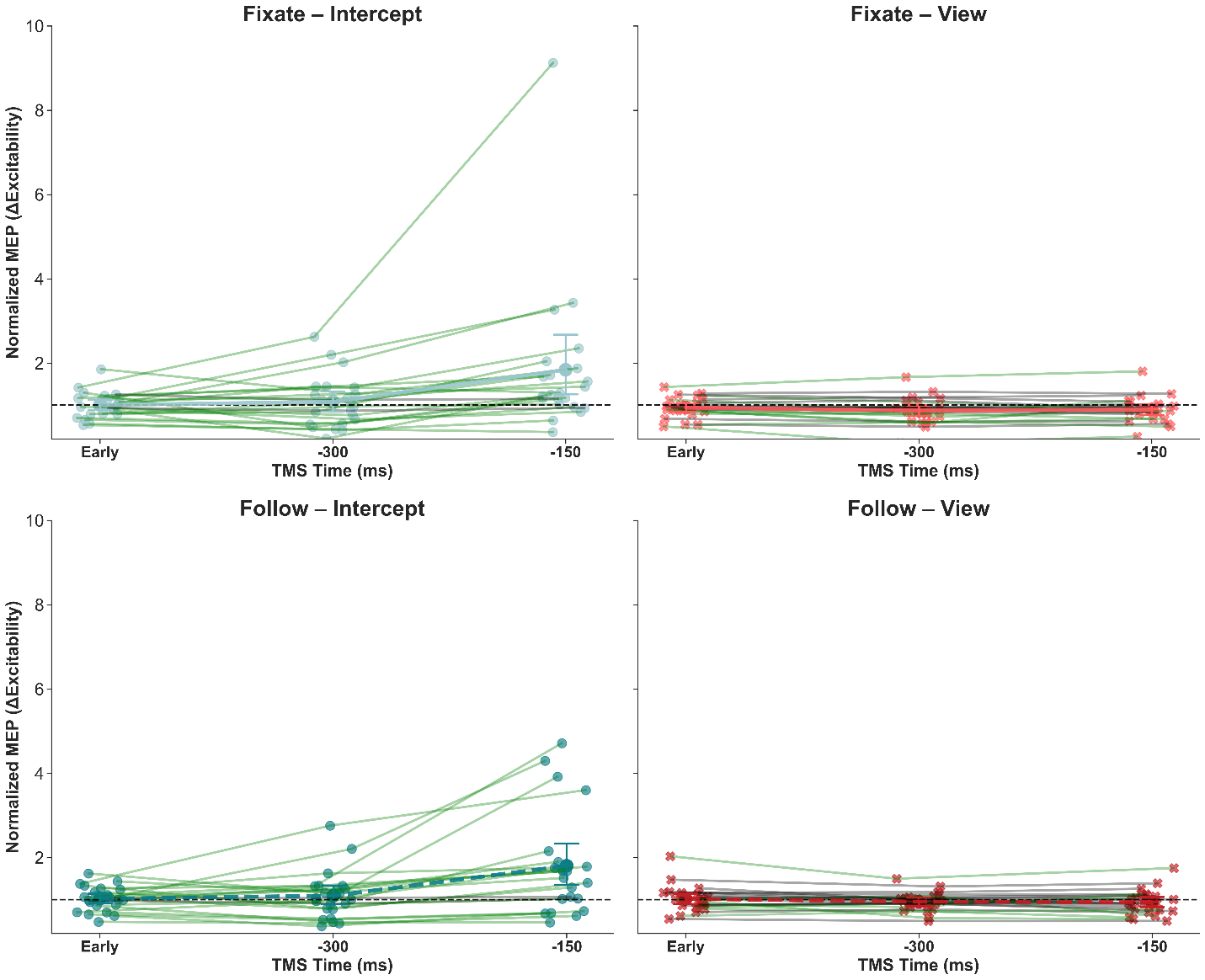

**Supplementary Figure 1.** Subject-level mean normalized MEP amplitudes are shown for each TMS time point (Early, -300 ms, -150 ms) separately for the four Task × Gaze conditions (Intercept-Fixate, Intercept-Follow, View-Fixate, View-Follow). Each point represents a participant’s mean normalized MEP at that TMS latency, overlaid on group means ± 95% CI to illustrate variability across individuals. This visualization was added in response to reviewer feedback highlighting the relatively large error bars in Figure 5, particularly for the Intercept conditions at –150 ms. Participant-level data indicate that the late increase in corticospinal excitability (CSE) during interception was not driven by only a small subset of individuals. Using a descriptive criterion (normalized MEP > 1.0 relative to baseline), facilitation at -150 ms was observed in 15/20 participants in the Intercept–Fixate condition, 15/20 in the Intercept-Follow condition, 6/20 in the View-Fixate condition, and 9/20 in the View-Follow condition. Thus, the larger error bars in the Intercept conditions primarily reflect variability in the magnitude of facilitation rather than its presence across participants.
